# Systematic In Silico Off-Target Assessment of siRNAs: Integrated Tissue-Specific Scoring and Cross-Species Preclinical Model Selection with TargetSureR

**DOI:** 10.64898/2026.05.28.728477

**Authors:** Shuai Ni, Kejia Kan, Fengjiao Zhu, Lina Wang, Wei Wang, Ning Wu

## Abstract

Small interfering RNAs (siRNAs) have become a transformative class of nucleic acid therapeutics for clinical disease treatment, yet sequence-dependent off-target silencing continues to pose a major safety barrier that hinders their preclinical refinement and large-scale translational application. Existing bioinformatics tools only support partial off-target evaluation, either focusing on basic sequence optimization or simple seed-region scanning, and fail to deliver systematic, multi-dimensional and reproducible safety assessment for siRNA lead screening. To fill this gap, we developed TargetSureR, a lightweight, modular and CRAN-compatible open-source R package dedicated to full-process siRNA off-target risk profiling. This tool accommodates dual sequence-based and precomputed position-based inputs, integrates GTEx multi-tissue expression data and curated cancer, adverse-event and immune gene panels, and establishes a seven-dimensional scoring framework to stratify off-target risks into four hierarchical tiers. It further enables tissue-specific safety characterization and quantitative cross-species model selection, with an Ensembl API fallback mechanism ensuring high transcript annotation resolution. Built purely in R with no external shell dependencies, TargetSureR provides a standardized, robust and user-friendly workflow for systematic siRNA preclinical safety evaluation, and is freely available at https://github.com/nishuai/TargetSureR.

## Introduction

Two decades after the discovery of RNA interference (RNAi)-mediated sequence-specific gene silencing in mammalian cells, RNAi has evolved from a basic laboratory technique into a mature therapeutic modality. The 2018 FDA approval of patisiran, a lipid nanoparticle-formulated siRNA drug for hereditary transthyretin amyloidosis (hATTR), represented a pivotal breakthrough, achieving durable clinical efficacy in previously untreatable disease [2,3,4]. Together with the concurrently approved antisense drug inotersen for the same indication [15], this milestone established nucleic acid therapeutics as a distinct drug class. Subsequent successive approvals of multiple RNAi drugs have validated the expanding clinical potential of this technology across hepatic, renal, ocular and neurological disease indications, consistent with prior clinical pipeline forecasts [1]. Advanced chemical modifications and GalNAc conjugation strategies have largely resolved delivery bottlenecks [5], shifting the core safety focus to siRNA sequence-dependent off-target risks.

Despite precise Watson–Crick base pairing, siRNAs exhibit widespread off-target effects with bimodal mechanisms. The siRNA guide strand mimics endogenous microRNA function, triggering pervasive off-target silencing via seed-region matching to 3’UTR partial complementary sites [6,7]. In addition, full or near-full complementarity binding in coding regions (CDS) induces Argonaute-2-mediated direct mRNA cleavage, a high-risk event neglected by conventional seed-based evaluation methods. These dose-dependent and reproducible off-target alterations disrupt genetic screening reliability, interfere with pharmacological effects and induce*in vivo* toxicity [6,7]. As RNAi therapeutics transition from short-term experimental knockdown to long-term clinical administration, regulatory requirements mandate comprehensive, human tissue-relevant characterization of full-spectrum sequence-derived off-target risks prior to clinical trials, which cannot be fully fulfilled by current computational tools.

Existing siRNA safety analysis tools suffer from severe functional fragmentation. Traditional design tools only optimize siRNA sequences by filtering high-risk seed matches at the design stage, lacking systematic profiling for finalized or chemically modified candidates. Tools such as SeedMatchR enable seed-mediated off-target detection but rely on experimental RNA-seq data and exclusively cover 3’UTR seed events, ignoring pathogenic CDS full-complementarity off-targets. Beyond incomplete site detection, standardized translational safety assessment requires multi-dimensional verification, including transcript regional classification, oncogene and immune/adverse-event risk gene annotation [13,14], GTEx-based multi-tissue expression profiling [12], and cross-species conservation evaluation for preclinical model validation. Although supporting public omics resources are available, their customized integration requires independent scripting, causing inconsistent analytical standards and poor reproducibility that biases candidate screening and hinders RNAi translational research.

To address these limitations, we developed TargetSureR, a reproducible, integrated R package for end-to-end siRNA off-target safety evaluation. It supports two mainstream input modes (raw siRNA sequences and precomputed positional data) with unified standardized outputs. The tool enables one-stop enrichment annotation covering transcript location classification, risk gene labeling and multi-tissue expression quantification. Its customizable seven-dimensional composite scoring system stratifies siRNA off-target risks into four grades for quantitative candidate screening. It also integrates cross-species ortholog analysis across six common preclinical species to guide rational animal model selection. As a lightweight, CRAN-compatible open-source pipeline, TargetSureR uniquely unifies dual-mode off-target identification, multi-omics risk annotation, tissue-level safety assessment, quantitative risk grading and cross-species validation. This study systematically describes its architecture, validates its performance, compares it with existing tools, and prospects future optimizations, aiming to provide a standardized and user-friendly platform for preclinical safety evaluation and lead optimization of RNAi therapeutics.

## Materials and Methods

### Package architecture and dependencies

TargetSureR is implemented as an R package (≥4.1.0) released under the MIT licence. The package is organised as a thin orchestration layer over a small number of well-characterised Bioconductor and CRAN dependencies, so that no shell-level binaries (BLAST, bowtie, RNAhybrid, etc.) need to be installed alongside it. Hard dependencies are limited to dplyr, tidyr, tibble, and readr for tabular manipulation. Annotation, sequence, and tissue-expression dependencies — SeedMatchR [10], ensembldb [11], Biostrings, AnnotationFilter, AnnotationHub, and GenomicRanges — are declared under the Suggests: field and loaded on demand, so users who only need one input mode (Section 2.2) do not pay the cost of installing the other. Built-in reference data are kept under 10 KB in total to comply with CRAN guidelines, with full-scale GTEx and ortholog matrices supplied at runtime by the user.

### Per-site rule-based risk score

Each off-target site is assigned an integer risk score ranging from 1 to 10 by the compute_risk_score() function, which quantifies risk by integrating the transcript region type and siRNA match specificity. The scoring framework adopts a region-based base score and a match-quality bonus score. Specifically, canonical CDS binding sites receive a base score of 5, 3’UTR sites are assigned a base score of 3, and all other sites located in 5’UTR or non-coding RNA regions are given a base score of 1. An additional bonus score of 2 is added for full-sequence complementary binding events, while partial matches carry no bonus. The final risk score is defined as the sum of the base score and bonus score, capped at a maximum value of 10. This rule-based scoring system follows well-established biological mechanisms. Full CDS complementarity triggers direct mRNA cleavage with high toxic risk [6], while 3’UTR seed matching represents the most prevalent off-target pattern in RNAi experiments [6,7]. In contrast, hits in 5’UTR and non-coding regions exert marginal and unstable effects. The parameter-free scoring strategy ensures transparent, reproducible and model-independent risk evaluation for all off-target sites.

### Biological enrichment

enrich_annotations() appends gene classification labels and tissue expression data to the core off-target table. The package includes three curated gene sets: adverse-event genes, an enhanced cancer gene panel covering oncogenes and tumour suppressors, and ImmPort-derived immune genes [13,14]. Gene matching is case-insensitive, and users may replace default panels with custom gene lists for specific research needs. Tissue expression data are imported as a matrix with gene symbols and tissue-specific expression values. A sample dataset is provided for reference, while the full GTEx v10 median-TPM matrix covering 59,033 genes across 68 tissues is recommended for formal analysis [12]. Genes missing expression data are assigned a value of zero. The final enriched dataset expands the original 10 columns to 81 columns with additional gene labels and tissue expression values.

### Per-siRNA composite score and risk tiers

The rank_sirna() function aggregates site-level off-target annotations to generate a unified risk profile for each siRNA, integrating seven independent metrics that quantify different types of off-target burden, with corresponding default weights listed in the following table.

All metric weights can be customized and are automatically normalized to a total value of one. Raw measurements are scaled linearly from 0 to 1 using either the built-in 94-siRNA reference cohort or the input dataset. Cohort-based normalization produces stable absolute risk scores for individual siRNAs, whereas self-normalization applies only to batch analyses. The final composite score is calculated as the weighted sum of normalized metrics, with lower values representing safer siRNAs. All sequences are ranked and stratified into four customizable risk tiers (Low, Medium, High, Critical), and candidate siRNAs can be explicitly labelled for easy identification.

This multi-metric system differentiates off-target events with distinct molecular mechanisms. The highest weight is assigned to high-risk hits in cancer, immune and adverse-event-associated genes, which dominate toxic and regulatory-relevant off-target effects. Additional counting metrics prevent score bias when severe off-target events are absent, while separate terms quantify CDS full-complementarity cleavage and 3’UTR seed-mediated repression, which respond differently to chemical modification strategies [5]. The original minimum free energy metric was discontinued, with its weight reassigned to high-risk gene hits, removing dependencies on external tools and ensuring consistent scoring for both position-based and sequence-based analysis modes.

### Per-gene tissue-attention score

tissue_safety() performs gene-level risk evaluation by integrating four key indicators: primary tissue expression level, maximum site risk score, proportion of high-risk CDS full-complementarity hits, and key risk gene attribution. Each indicator is normalized and weighted to generate a unified bio_attention score, where tissue expression occupies the dominant weight, emphasizing that only off-target genes expressed in drug-targeted tissues contribute to clinical risk. Genes are ranked by this score to screen priority safety-risk genes for further validation.

### Cross-species animal-model recommendation

The species_match() function evaluates whether preclinical animal models recapitulate human off-target profiles. It queries ortholog information and sequence identity for each off-target gene from an orthology table, and calculates conservation scores by ortholog identity: 1.0 for identity ≥90%, 0.6 for 80%–89%, 0.3 for identity <80%, and 0 if no ortholog exists. The analysis targets critical risk genes by default. For each siRNA, species-specific scores are averaged across relevant sites, with outputs including scoring matrices, site details and model recommendations. It supports six widely used species in RNAi drug development and allows custom species selection. This orthology-based approach runs efficiently without genome assemblies, avoiding heavy computation from full cross-species alignment. A supplementary seed-matching module will be added in future versions to improve assessment for siRNAs with few critical off-target genes.

### Dataset curation and evaluation cohort

The seven-dimensional composite scoring system of TargetSureR was normalized using a rigorously curated reference cohort derived from the public MIT/ICBP siRNA Database [16,17,18,19], a well-established resource for annotated RNAi reagents with documented gene information and experimental profiles. A total of 144 original entries were initially retrieved, and 50 long hairpin shRNA constructs were excluded to retain 94 valid siRNAs targeting 66 unique human genes (Table 2). Most entries contained standard RefSeq accessions and complete strand information, supporting reliable and standardized downstream calibration. For end-to-end tool validation, a 97-siRNA evaluation cohort was constructed, combining three distinct KRAS-targeting siRNAs and the 94 curated reference siRNAs. All sequences were analyzed using the complete Mode A pipeline incorporating dual seed/full-length matching, GTEx-based tissue expression profiling [12], curated risk gene annotation [13,14], and reference-cohort normalization, with fixed analytical parameters to guarantee stable, reproducible and unbiased tool performance assessment.

**Table 1.**
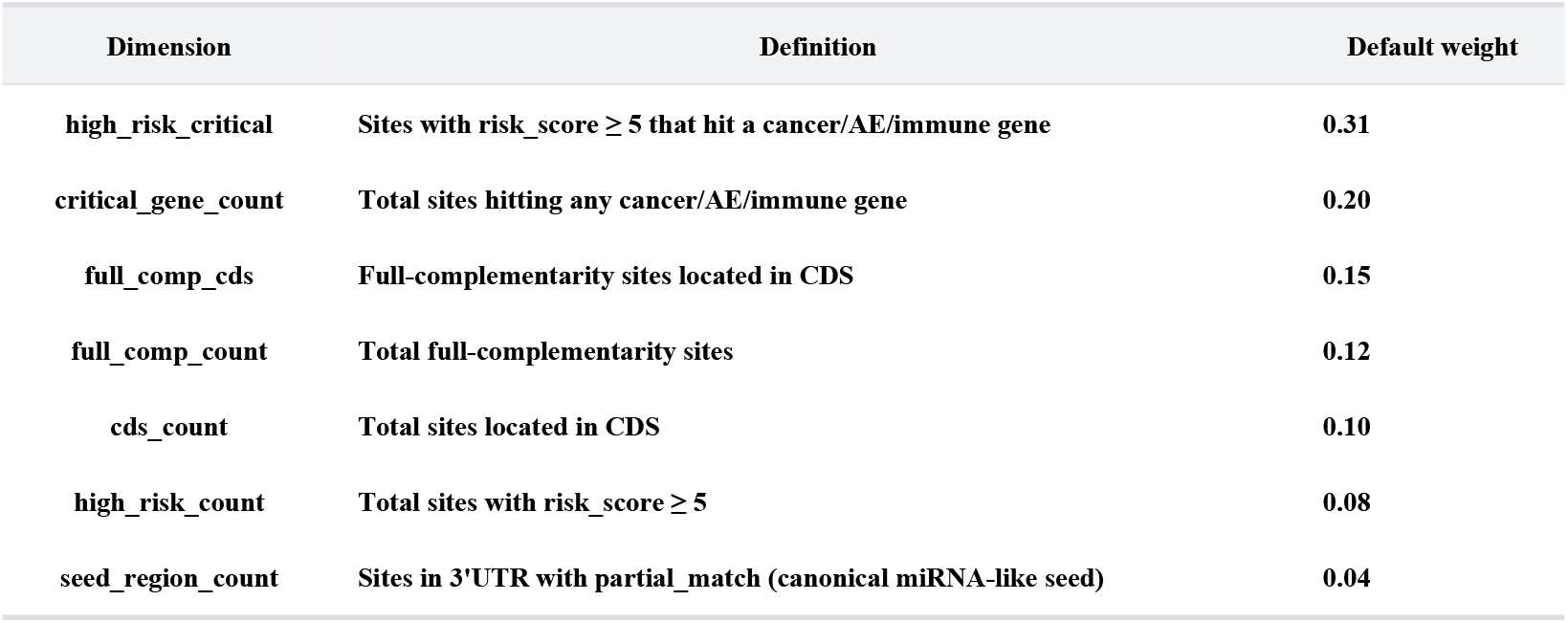
Seven-dimensional scoring metrics and default weight configuration of the rank_sirna() function.

**Table 2.** Summary statistics for the MIT/ICBP siRNA cohort curation.

| Item | Count |
| --- | --- |
| Entries retrieved from MIT/ICBP database | 144 |
| Long hairpin/shRNA entries excluded (>30 nt) | 50 |
| siRNA entries retained (19–30 nt) | 94 |
| Unique gene symbols | 66 |
| Entries with RefSeq accession | 89 |
| Entries with both sense and antisense strands | 4 |

## Results

We applied TargetSureR end-to-end to the 97-siRNA evaluation cohort described in Section 2.10 — three KRAS-targeting query siRNAs (KRAS_si8, KRAS_si11, KRAS_si17 from Ramalingam and Arumugam [20]) compared against the 94-siRNA MIT/ICBP reference cohort that ships with the package as the built-in reference_set (Section 2.9). Mode A scanned each query against three feature classes (3’UTR, 5’UTR, CDS) of the human protein-coding transcriptome (Ensembl release 113, ∼19,000 transcripts per class after longest-per-gene reduction), yielding **19**,**224 off-target sites** in total across the three guides (KRAS_si8 2,588 sites; KRAS_si17 5,675; KRAS_si11 10,961). All numerical results below are derived from one end-to-end run of paper/realrun/demo_3kras_vs_mit94.R; figures referenced are the unmodified output of generate_figures(). The 94-siRNA reference distributions used for normalisation and overlay are the corresponding fields of the reference_set data object distributed with the package.

### Composite ranking places two of three KRAS guides in the Low tier

Composite scaling against the per-dimension minima and maxima of the 94-siRNA MIT/ICBP cohort (built-in tier_thresholds, Section 2.9) places KRAS_si17 and KRAS_si8 in the Low tier and KRAS_si11 in the Medium tier (Table 3; Figure 1). All three queries fall on the safer side of the reference distribution: the 94-siRNA reference cohort has a 25th-percentile composite score of 0.534 and a median of 0.583, whereas KRAS_si17 (0.199), KRAS_si8 (0.216), and KRAS_si11 (0.431) sit at, well below, and just below the 25th percentile of the reference cohort, respectively. By construction the reference cohort is concentrated in the High tier (79 of 94 reference siRNAs, 84 %), with 14 in Medium, one in Low and zero in Critical, so any query reaching the Low tier is comparatively clean.

**Table 3.** Per-siRNA composite metrics for the three KRAS query siRNAs, scored under default seven-dimensional weights and normalisation against the 94-siRNA MIT/ICBP cohort.

| siRNA | Rank | Total off-targets | High-risk × critical | Critical-gene hits | Full-comp (CDS) | Full-comp (total) | CDS hits | High-risk hits | Seed (3'UTR) | Composite | Tier |
| --- | --- | --- | --- | --- | --- | --- | --- | --- | --- | --- | --- |
| KRAS_si17 | 1 | 5,675 | 19 | 47 | 0 | 1 | 1,058 | 1,059 | 4,351 | 0.199 | Low |
| KRAS_si8 | 2 | 2,588 | 12 | 27 | 1 | 1 | 1,125 | 1,125 | 1,338 | 0.216 | Low |
| KRAS_si11 | 3 | 10,961 | 40 | 86 | 1 | 1 | 5,083 | 5,083 | 5,271 | 0.431 | Medium |

For comparison, the 94-siRNA MIT/ICBP reference cohort has a median total-off-target count of 5,316 (interquartile range 3,298–7,408), a median high_risk_critical of 14 (IQR 9–21) and a median critical_gene_count of 33.5 (IQR 23.3–52.8). All three KRAS queries are at or below the cohort medians on these dimensions — KRAS_si8 notably sits below the 25th percentile of the reference on every count except cds_count/high_risk_count — and the ranking violin (Figure 1) shows the three coloured diamonds anchoring the lower tail of the reference distribution.

**Figure 1.**
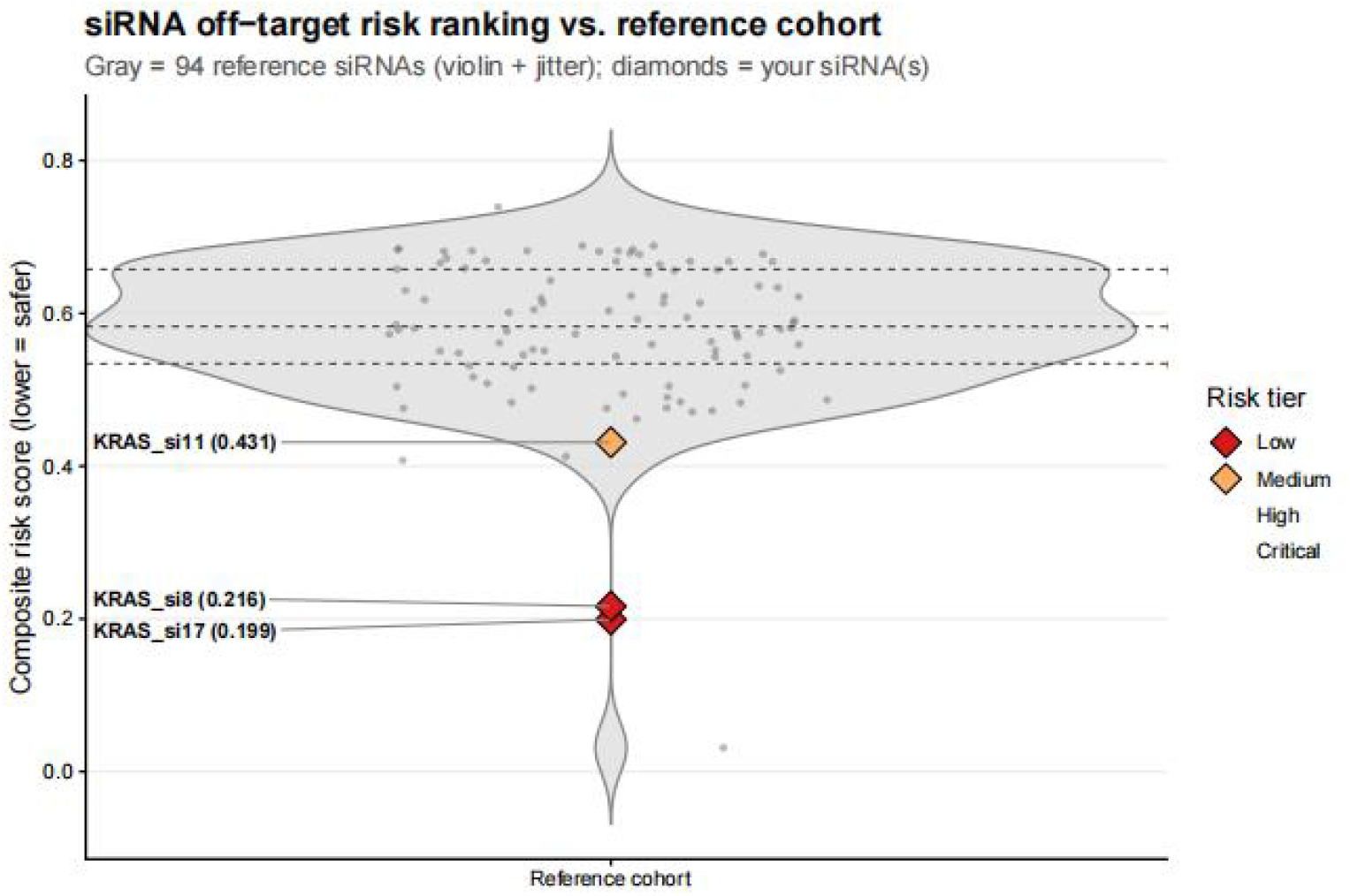
Composite risk ranking of three KRAS-targeting siRNAs normalized against the MIT/ICBP reference cohort. Violin plot illustrating the distribution of composite risk scores across the 94-siRNA MIT/ICBP reference cohort, with overlaid individual scores of three tested KRAS siRNA candidates. The reference cohort exhibits a median composite score of 0.583 and a 25th percentile value of 0.534, with most reference siRNAs (84 %) classified into the High risk tier. The colored diamond markers represent KRAS_si17 (0.199, Low tier), KRAS_si8 (0.216, Low tier), and KRAS_si11 (0.431, Medium tier), all of which rank below the 25th percentile of the reference distribution and demonstrate superior safety profiles compared with most canonical siRNA reagents in the built-in dataset.

### Per-dimension decomposition reveals distinct off-target fingerprints

Although the three guides share a target gene, their off-target landscapes differ substantially in shape. The KRAS_si8 profile is dominated by CDS-localised hits (1,125 of 2,588 sites, 43 %; with one full-complementarity match in CDS) and shows a comparatively sparse 3’UTR seed signature (1,338 sites). KRAS_si17 distributes its hits in the opposite direction: only 1,058 of 5,675 sites (19 %) fall in CDS, while 4,351 sites (77 %) are 3’UTR seed-region matches, indicating a guide whose seed sequence is over-represented in the human 3’UTR-ome but whose CDS-cleavage liability is small. KRAS_si11 shows both modes elevated together (5,083 CDS sites and 5,271 3’UTR seed sites), giving it the heaviest aggregate burden of the three and pushing it to the Medium tier despite having only one full-complementarity hit.

The risk-distribution stacked bar (Figure 2) places these three profiles inside the cohort distribution: KRAS_si8 sits at the low-burden end of the cohort, KRAS_si17 in the lower middle, and KRAS_si11 near the median. The radar-style decomposition shows that no query reaches the cohort 75th percentile on any of the seven dimensions — consistent with their Low/Low/Medium tier assignment.

**Figure 2.**
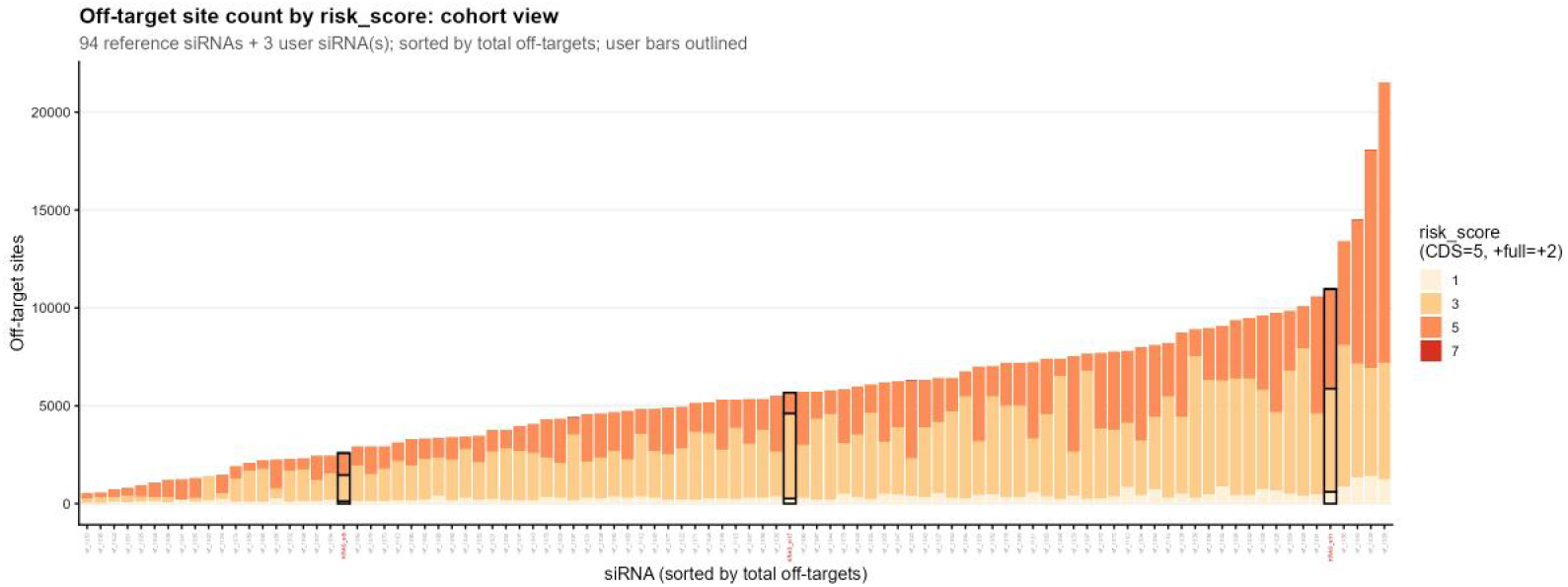
Divergent off-target landscape profiles of three KRAS-targeting siRNAs across transcript regions. Stacked bar plot illustrating the decomposed off-target distributions of KRAS_si8, KRAS_si17, and KRAS_si11 across CDS and 3’UTR regions, benchmarked against the 94-siRNA MIT/ICBP reference cohort. KRAS_si8 exhibits CDS-dominant off-target signatures with limited 3’UTR seed hits, whileKRAS_si17 presents predominantly 3’UTR seed-mediated off-target events with minimal CDS cleavage risk. In contrast, KRAS_si11 displays substantially elevated off-target burdens in both regions, resulting in the highest overall risk and Medium-tier classification. All three candidates remain below the 75th percentile of the reference cohort across all seven evaluation dimensions, consistent with their assigned low-to-moderate risk stratification.

### Per-gene attention identifies cancer-panel hits as the actionable shortlist

Aggregating sites to genes throughtissue_safety() reduces 19,224 site-level entries to a ranked list of unique off-target genes. Because all three queries target KRAS, the top gene of the attention table is **KRAS itself** with a bio_attention of 0.682 (9 hits, max risk score 7, two of which are full-complementarity in CDS, on-target by design). The next nine highest-attention genes are all members of the curated cancer or adverse-event panels (Table 4) and represent the project-actionable shortlist: each one combines measurable expression in the assumed primary tissue, multiple sites across the three guides, and membership in at least one critical-gene panel.

**Table 4.** Top ten off-target genes by bio_attention (excluding on-target). All entries hit at least one of the cancer, adverse-event, or immune panels.

| Rank | Gene | n hits | primary TPM | max risk | bio_attention | Notes |
| --- | --- | --- | --- | --- | --- | --- |
| 1 | NRAS | 1 | 26.75 | 5 | 0.667 | RAS-family paralogue, oncogene |
| 2 | EGFR | 2 | 12.32 | 5 | 0.478 | Cancer + AE |
| 3 | PIK3CA | 6 | 10.18 | 5 | 0.450 | Cancer |
| 4 | MSH6 | 4 | 10.02 | 5 | 0.448 | AE (Lynch syndrome) |
| 5 | BRCA1 | 3 | 5.13 | 5 | 0.384 | Cancer + AE |
| 6 | PALB2 | 3 | 5.12 | 5 | 0.384 | Cancer + AE |
| 7 | TP53 | 2 | 4.07 | 5 | 0.370 | Cancer + AE |
| 8 | CTNNB1 | 1 | 3.91 | 5 | 0.368 | Cancer |
| 9 | MYC | 1 | 2.47 | 5 | 0.349 | Cancer |
| 10 | PTEN | 5 | 2.05 | 5 | 0.343 | Cancer + AE |

Of immediate concern is the recurring hit on **NRAS**, the closest paralogue of the on-target gene, which combines high expression (TPM 26.8) with a partial-match seed-region match — a classic seed-mediated cross-silencing risk for any guide directed against a RAS-family member. The second-tier attention list (PIK3CA, MSH6, BRCA1, PALB2, TP53, PTEN) is dominated by tumour suppressors and DNA-repair genes whose silencing would be consequential for any chronic-dosing oncology candidate. The gene × siRNA bubble plot (Figure3 ) shows that these high-attention genes are not uniformly hit across the three queries: the heaviest critical-gene burden falls on KRAS_si11 (86 critical-gene hits), confirming the per-dimension result that this guide concentrates the panel-relevant risk among the three.

**Figure 3.**
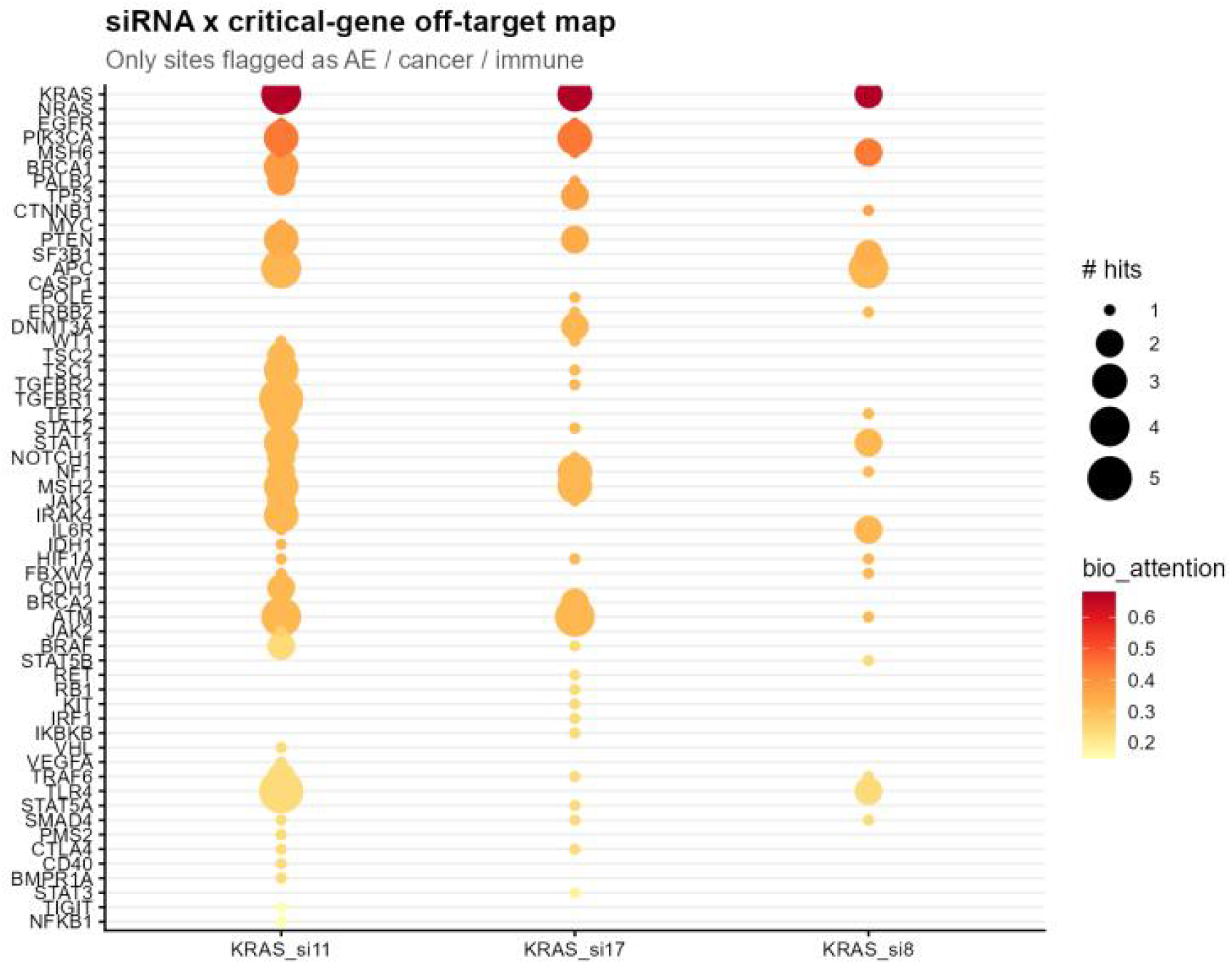
Gene-level off-target attention profiling and high-risk gene distribution of three KRAS-targeting siRNAs. Gene–siRNA bubble plot visualizes the screened high-risk off-target genes ranked by bio_attention scores, which integrate off-target hit counts, tissue-specific expression levels, and maximum risk scores. Excluding the on-target KRAS gene, the top-ranked off-target candidates are predominantly well-characterized oncogenes, tumour suppressors, and disease-relevant adverse-event genes. The plot demonstrates uneven risk distribution across the three siRNA guides, with KRAS_si11 producing the highest burden of critical-gene off-target hits. Notably, the highly expressed RAS paralogue NRAS represents a prominent seed-mediated cross-silencing risk, while other enriched genes involved in DNA repair and tumour regulation indicate substantial safety concerns for long-term therapeutic administration.

### Animal-model recommendation: cynomolgus is the consistent first choice

Cross-species ortholog conservation across the three KRAS guides on critical-gene off-targets returns the per-species score matrix in Table 5 (Figure 4; Figure 5).

**Table 5.** Layer-1 ortholog conservation score on critical off-target genes (range 0–1; higher = better animal model).

| siRNA | mouse | rat | cyno | rhesus | rabbit | dog | Recommended |
| --- | --- | --- | --- | --- | --- | --- | --- |
| KRAS_si8 | 0.259 | 0.193 | 0.407 | 0.348 | 0.289 | 0.230 | cyno |
| KRAS_si17 | 0.153 | 0.183 | 0.277 | 0.268 | 0.211 | 0.194 | cyno |
| KRAS_si11 | 0.164 | 0.145 | 0.281 | 0.272 | 0.177 | 0.200 | cyno |

**Figure 4.**
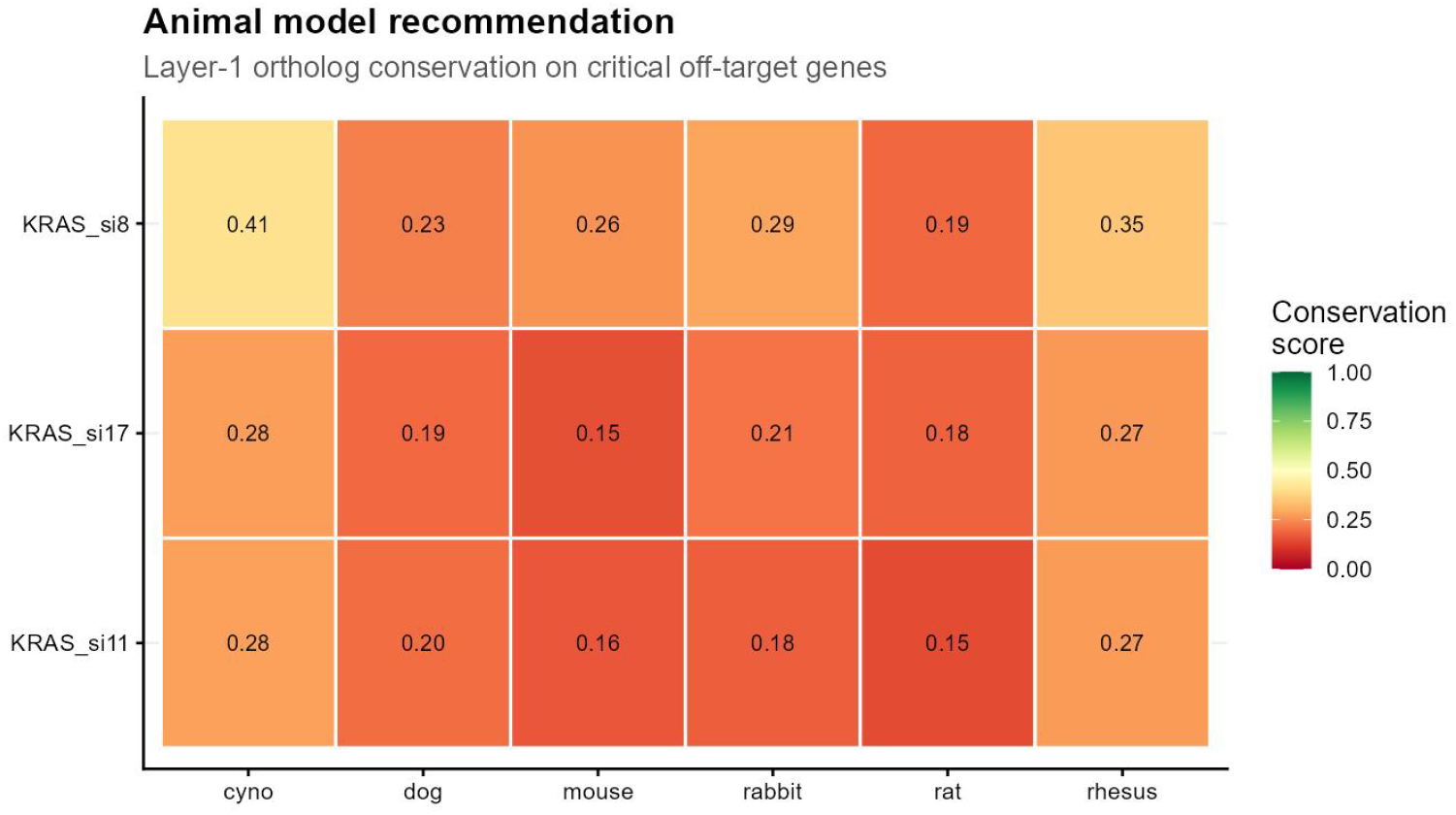
Cross-species ortholog conservation heatmap of critical off-target genes for three KRAS-targeting siRNAs. Heatmap visualization of standardized Layer-1 ortholog conservation scores (range: 0–1) across six common preclinical species, including mouse, rat, cynomolgus monkey, rhesus monkey, rabbit, and dog. Higher scores indicate higher similarity to human off-target profiles. Consistent across all three siRNA candidates, cynomolgus monkey yields the highest conservation values, followed by rhesus monkey, while rodent models exhibit the lowest genetic consistency. This result supports the preferential selection of non-human primates for preclinical safety evaluation of KRAS-targeted siRNA therapeutics.

Cynomolgus monkey (*Macaca fascicularis*) is the top recommended model for all three queries, with rhesus monkey a close second and rodents trailing. This ordering is consistent with the 94-siRNA reference cohort (cyno median 0.260, rhesus 0.242, rabbit 0.180, dog 0.160, mouse 0.160, rat 0.150), confirming that for a cancer-relevant target gene with a critical-gene off-target panel dominated by tumour suppressors and oncogenes, the non-human-primate models reproduce the human off-target profile substantially more faithfully than the rodent or non-primate alternatives. The percentile bar plot (Figure 04c_species_percentile) places KRAS_si8’s cynomolgus score at the upper end of the reference distribution, and Figure 5 shows the three queries’ diamonds tracking the reference-cohort violin pattern across all six species.

**Figure 5.**
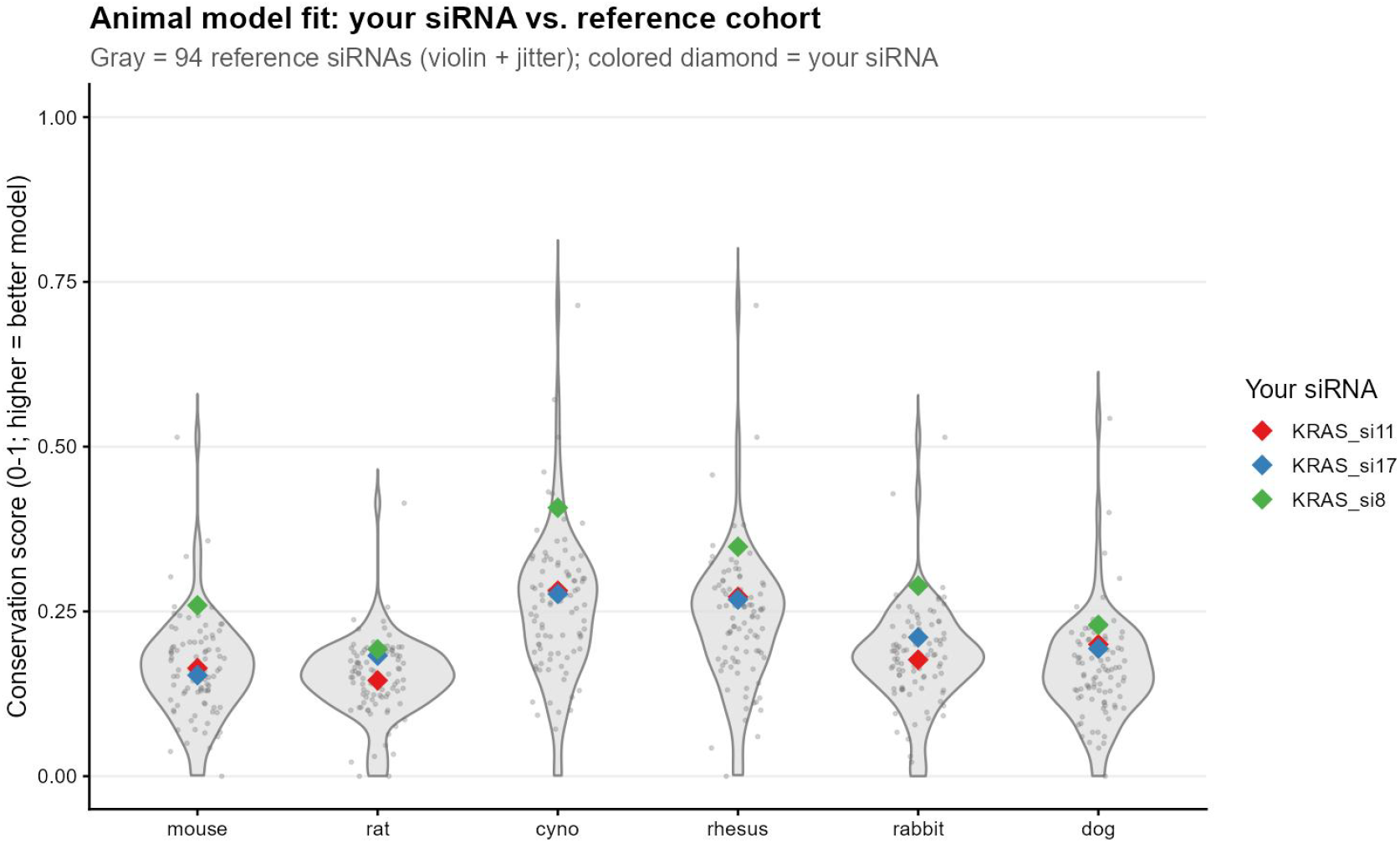
Cross-species conservation performance of KRAS siRNAs relative to the MIT/ICBP reference cohort. Violin and percentile distribution plots benchmarking species conservation scores of the three KRAS-targeting siRNAs against the 94-siRNA reference cohort. The overall species preference trend of the tested siRNAs aligns with the reference baseline, with non-human primates outperforming rodents and other non-primate species. In particular, KRAS_si8 achieves a superior cynomolgus conservation score located at the upper tail of the reference distribution, demonstrating robust human-mimicking off-target profiling of primate models for oncogenic target genes.

### Summary of the evaluation

Three observations from this evaluation are directly relevant to the use of TargetSureR in lead selection. **First**, the seven-dimensional composite separates structurally similar candidates: although KRAS_si8, KRAS_si11 and KRAS_si17 all target the same mRNA and were short-listed together by upstream rule-based filters [20], TargetSureR places them across two risk tiers (KRAS_si8 and KRAS_si17 Low, KRAS_si11 Medium) on the basis of total off-target burden and critical-gene engagement, with KRAS_si11 carrying ∼4× the off-target load and ∼3× the critical-gene engagement of KRAS_si8. **Second**, the per-gene bio_attention shortlist concentrates roughly 19,000 sites into a ranked panel of off-target genes, the top ten of which are all curated cancer or adverse-event genes. The recurring partial-match hit on the RAS paralogue NRAS — driven by seed-region complementarity and not by full-length identity — is exactly the class of liability that seed-only design tools (siDirect [9]) and post-treatment RNA-seq scanners (SeedMatchR [10]) cannot jointly surface, and that TargetSureR’s combined region × match-type × tissue-expression × gene-list view does. **Third**, the animal-model recommender unambiguously favours non-human primates for this target class, with cynomolgus monkey scoring 0.28–0.41 across all three guides versus 0.15–0.26 for rodents.

We make no claim of absolute predictive accuracy from this single evaluation; absent a held-out experimental dataset, the analysis demonstrates only that TargetSureR’s composite ranking, per-gene attention shortlist, and species recommender are internally consistent and produce mechanistically interpretable distinctions between three published candidates against a real 94-siRNA reference background. Section 4 places TargetSureR in the context of existing tools and discusses limitations of the current implementation.

### Benchmark against existing tools and discussion of limitations

Table 6 systematically compares TargetSureR with mainstream published siRNA design and off-target prediction tools. The evaluation covers seven core dimensions that determine tool suitability for regulatory-level lead screening, including dual scanning mechanisms (seed-based and full-complementarity matching), tissue expression filtering, curated risk gene annotation, cross-species model recommendation, precomputed positional data compatibility, pure offline R execution, and ongoing tool maintenance status.

**Table 6.** Feature comparison between TargetSureR and published siRNA off-target or design tools. ✓ = implemented; ✗ = not implemented; (✓) = partially or indirectly supported.

| Tool | Year | Seed scan | Full-comp scan | Tissue expression filter | Cross-species recommendation | Reference Cohort | Standalone / pure R | Active availability |
| --- | --- | --- | --- | --- | --- | --- | --- | --- |
| TargetSureR | 2026 | ✓ | ✓ | ✓ (GTEx v10, 68 tissues) | ✓ (6 species) | ✓ | R | ✓ |
| siDirect 2.0 [9] | 2009 | ✓ | ✗ | ✗ | ✗ | ✗ | Web | (✓) |
| siDirect 1.0 [8] | 2004 | ✓ | ✗ | ✗ | ✗ | ✗ | Web | ✗ |
| SeedMatchR [10] | 2024 | ✓ | ✗ | ✗ | ✗ | ✗ | R | ✓ |
| siSPOTR [21] | 2013 | ✓ | ✗ | ✗ | ✗ | ✗ | Web | ✗ |
| OligoWalk [22] | 2008 | ✗ | ✓ | ✗ | ✗ | ✗ | Web | ✗ |
| Das et al. tool [23] | 2013 | ✓ | ✗ | ✗ | ✗ | ✗ | Web | ✗ |

TargetSureR outperforms existing tools via five unique functionalities. It integrates tissue-specific expression filtering to prioritize actionable off-target risks and annotates clinically relevant cancer, adverse-event and immune genes for regulatory-oriented risk scoring. It supports flexible dual-input modes and establishes standardized multi-tier risk stratification based on reference-cohort normalization. It also provides quantitative cross-species conservation analysis to guide preclinical animal model selection for translational research.

## Discussion

While TargetSureR exhibits comprehensive advantages over existing siRNA off-target evaluation tools in multi-dimensional annotation, quantitative risk grading and cross-species translational assessment, the current platform still has several technical limitations that can be further optimized in future iterations. First, the normalization and risk-tier partitioning of the built-in reference_set are calibrated based on 94 MIT/ICBP-derived laboratory siRNAs targeting 66 heterogeneous endogenous genes. This reference dataset primarily covers conventional unmodified siRNA reagents, which may not fully represent the characteristics of clinical-grade chemically modified siRNAs, such as GalNAc-conjugated candidates. Therefore, users conducting hepatic or ocular RNAi therapeutic development are recommended to construct project-specific custom reference cohorts to achieve more accurate, target-adaptive risk stratification.

Second, the current seven-dimensional composite scoring system is a rule-based model dependent on transcript location and match type (Section 2.3), without incorporating RNA duplex thermodynamic stability parameters. This design avoids reliance on non-CRAN external tools including RNAhybrid and RNAfold, ensuring pure-R offline operability and environment independence. However, this strategy results in identical risk scoring for off-target sites with consistent regional and matching features but distinct thermodynamic stability, potentially overestimating the risk of low-stability full-complementary sites and underestimating the off-target potency of thermodynamically stable seed-mediated partial matches. Future updates will introduce R-based stability proxies or lightweight interfaces to professional thermodynamic tools to refine scoring accuracy.

Third, the sequence-based scanning pipeline of Mode A has limitations in transcript regional annotation. The annotate_from_sequence() function performs separate scanning for 3’UTR, 5’UTR and CDS features and assigns region labels based on single matched feature classes, leading to potential annotation ambiguity for sites spanning multiple transcript isoforms with overlapping regional attributes. In comparison, Mode B enables precise and unambiguous regional classification through quantitative calculation of mature mRNA structural lengths via ensembldb. Subsequent versions will unify Mode A with this isoform-aware annotation strategy to eliminate classification discrepancies between the two input modes.

Fourth, the package’s embedded ortholog_table_example is a simplified dataset designed only for tutorial and demonstration purposes, which is insufficient for formal preclinical translational analysis. Rigorous cross-species safety evaluation requires a full-scale ortholog matrix covering all human protein-coding genes, derived from Ensembl BioMart across six preclinical species. Users can generate customized complete ortholog resources following the open-source template scripts provided in the package to support production-grade analysis.

Fifth, the current algorithm lacks awareness of siRNA chemical modifications. All guide sequences are treated as unmodified 21-mer RNAs during risk calculation, ignoring the functional alterations brought by mainstream chemical optimizations. Common 2’-fluoro and 2’-O-methyl modifications profoundly modulate seed-region duplex stability and Argonaute loading kinetics; notably, seed-region 2’-O-methyl modification can reduce off-target silencing efficiency by approximately 10-fold [24]. Future optimization will integrate position-specific chemical modification parameters to establish a modification-adaptive scoring system tailored for clinical oligonucleotide candidates.

Finally, TargetSureR functions as a pure in silico predictive tool and does not support integration with post-treatment experimental transcriptomic data. In contrast, SeedMatchR [10] is specialized in verifying actual off-target alterations via RNA-seq differential enrichment analysis. The two tools are highly complementary: TargetSureR comprehensively predicts theoretical off-target risks at the pre-experimental stage, while SeedMatchR validates practical off-target effects in specific cell types and dosing contexts. Integrating RNA-seq-based enrichment verification into the TargetSureR pipeline will bridge in silico prediction and experimental validation, further improving the reliability and translational value of siRNA preclinical safety assessment.

## References

[1] Setten RL, Rossi JJ, Han SP. The current state and future directions of RNAi-based therapeutics. Nature Reviews Drug Discovery. 2019;18(6):421–446. doi:10.1038/s41573-019-0017-4. PMID:30846871.

[2] Adams D, Gonzalez-Duarte A, O’Riordan WD, Yang CC, Ueda M, Kristen AV, et al. Patisiran, an RNAi Therapeutic, for Hereditary Transthyretin Amyloidosis. New England Journal of Medicine. 2018;379(1):11–21. doi:10.1056/NEJMoa1716153. PMID:29972753.

[3] Hoy SM. Patisiran: First Global Approval. Drugs. 2018;78(15):1625–1631. doi:10.1007/s40265-018-0983-6. PMID:30251172.

[4] Kristen AV, Ajroud-Driss S, Conceição I, Gorevic P, Kyriakides T, Obici L. Patisiran, an RNAi therapeutic for the treatment of hereditary transthyretin-mediated amyloidosis. Neurodegenerative Disease Management. 2019;9(1):5–23. doi:10.2217/nmt-2018-0033. PMID:30480471.

[5] Khvorova A, Watts JK. The chemical evolution of oligonucleotide therapies of clinical utility. Nature Biotechnology. 2017;35(3):238–248. doi:10.1038/nbt.3765. PMID:28244990.

[6] Jackson AL, Linsley PS. Recognizing and avoiding siRNA off-target effects for target identification and therapeutic application. Nature Reviews Drug Discovery. 2010;9(1):57–67. doi:10.1038/nrd3010. PMID:20043028.

[7] Birmingham A, Anderson EM, Reynolds A, Ilsley-Tyree D, Leake D, Fedorov Y, et al. 3’ UTR seed matches, but not overall identity, are associated with RNAi off-targets. Nature Methods. 2006;3(3):199–204. PMID:16489337.

[8] Naito Y, Yamada T, Ui-Tei K, Morishita S, Saigo K. siDirect: highly effective, target-specific siRNA design software for mammalian RNA interference. Nucleic Acids Research. 2004;32(Web Server issue):W124–W129. PMID:15215364.

[9] Naito Y, Yoshimura J, Morishita S, Ui-Tei K. siDirect 2.0: updated software for designing functional siRNA with reduced seed-dependent off-target effect. BMC Bioinformatics. 2009;10:392. doi:10.1186/1471-2105-10-392. PMID:19948054.

[10] Cazares T, Higgs RE, Wang J, Ozer HG. SeedMatchR: identify off-target effects mediated by siRNA seed regions in RNA-seq experiments. Bioinformatics. 2024;40(1):btae011. doi:10.1093/bioinformatics/btae011. PMID:38192001.

[11] Rainer J, Gatto L, Weichenberger CX. ensembldb: an R package to create and use Ensembl-based annotation resources. Bioinformatics. 2019;35(17):3151–3153. doi:10.1093/bioinformatics/btz031. PMID:30689724.

[12] GTEx Consortium. The GTEx Consortium atlas of genetic regulatory effects across human tissues. Science. 2020;369(6509):1318–1330. doi:10.1126/science.aaz1776. PMID:32913098.

[13] Bhattacharya S, Andorf S, Gomes L, Dunn P, Schaefer H, Pontius J, et al. ImmPort: disseminating data to the public for the future of immunology. Immunological Research. 2014;58(2-3):234–239. doi:10.1007/s12026-014-8516-1. PMID:24791905.

[14] Bhattacharya S, Dunn P, Thomas CG, Smith B, Schaefer H, Chen J, et al. ImmPort, toward repurposing of open access immunological assay data for translational and clinical research. Scientific Data. 2018;5:180015. doi:10.1038/sdata.2018.15. PMID:29485622.

[15] Benson MD, Waddington-Cruz M, Berk JL, Polydefkis M, Dyck PJ, Wang AK, et al. Inotersen Treatment for Patients with Hereditary Transthyretin Amyloidosis. New England Journal of Medicine. 2018;379(1):22–31. doi:10.1056/NEJMoa1716793. PMID:29972757.

[16] MIT/ICBP siRNA Database. Sharp Laboratory, Koch Institute / MIT Center for Cancer Research, Massachusetts Institute of Technology. https://web.mit.edu/sirna/ (accessed 2026-05-15).

[17] Elbashir SM, Harborth J, Lendeckel W, Yalcin A, Weber K, Tuschl T. Duplexes of 21-nucleotide RNAs mediate RNA interference in cultured mammalian cells. Nature. 2001;411(6836):494–498. doi:10.1038/35078107. PMID:11373684.

[18] Doench JG, Petersen CP, Sharp PA. siRNAs can function as miRNAs. Genes & Development. 2003;17(4):438–442. doi:10.1101/gad.1064703. PMID:12600936.

[19] Reynolds A, Leake D, Boese Q, Scaringe S, Marshall WS, Khvorova A. Rational siRNA design for RNA interference. Nature Biotechnology. 2004;22(3):326–330. doi:10.1038/nbt936. PMID:14758366.

[20] Ramalingam PS, Arumugam S. Computational design and validation of effective siRNAs to silence oncogenic KRAS. 3 Biotech. 2023;13(11):350. doi:10.1007/s13205-023-03767-w. PMID:37780803. (PMC10541393.)

[21] Boudreau RL, Spengler RM, Hylock RH, Kusenda BJ, Davis HA, Eichmann DA, et al. siSPOTR: a tool for designing highly specific and potent siRNAs for human and mouse. Nucleic Acids Research. 2013;41(1):e9. doi:10.1093/nar/gks797. PMID:22941647.

[22] Lu ZJ, Mathews DH. OligoWalk: an online siRNA design tool utilizing hybridization thermodynamics. Nucleic Acids Research. 2008;36(Web Server issue):W104–W108. doi:10.1093/nar/gkn250. PMID:18490376.

[23] Das S, Ghosal S, Kozak K, Chakrabarti J. An siRNA designing tool with a unique functional off-target filtering approach. Journal of Biomolecular Structure and Dynamics. 2013;31(11):1343–1357. doi:10.1080/07391102.2012.736758. PMID:23140209.

[24] Jackson AL, Burchard J, Leake D, Reynolds A, Schelter J, Guo J, et al. Position-specific chemical modification of siRNAs reduces “off-target” transcript silencing. RNA. 2006;12(7):1197–1205. PMID:16682562.

